# Nucleobindin-1 *(Nucb1)* disruption affects feeding, metabolism, and glucose homeostasis in mice in an age-, sex-, diet- and light cycle-dependent manner

**DOI:** 10.1101/2024.03.07.584005

**Authors:** Atefeh Nasri, Emilio J Vélez, Jithine Jayakumar Rajeswari, Azadeh Hatef, Suraj Unniappan

## Abstract

**Background:** Nesfatin-1 (NESF-1), encoded in the calcium and DNA binding protein (Nucleobindin 2, NUCB2) is an orphan ligand with metabolic effects. Recently, our lab provided evidence for a NESF-1-like peptide (NLP) in a NUCB2-related precursor, NUCB1, in zebrafish and rodents. This research aims to determine whether endogenous NUCB1 is critical for energy homeostasis.

**Methods and Main Findings:** Global genetic disruption of *Nucb1* (*Nucb1* knockout/KO mice) led to increased food intake in chow-fed male and female mice across different points of light and dark phases. A similar increase in water intake was seen in female *Nucb1* KO mice but not in males. White adipose tissue weight was significantly increased in male and female *Nucb1* KO mice. Dark phase total activity was increased in male *Nucb1* KO mice, while it was decreased in female *Nucb1* KO mice compared to wildtype littermates. Energy derived from carbohydrates was raised during the dark phase; while energy derived from fat was significantly decreased in both male and female *Nucb1* KO mice. Male *Nucb1* KO mice were lighter in the early stages, but these differences disappeared as they aged. Meanwhile, no differences in bodyweight were observed in female *Nucb1* KO mice. Male *Nucb1* KO mice handled glucose better during an oral glucose tolerance test, while the opposite effect was found in an intraperitoneal (IP) glucose tolerance test. The above results from chow-fed mice were largely true in 10% and 60% fat diet-fed mice. A significant two-way interaction between mice group and time was observed on weekly food intake of male and female *Nucb1* KO mice fed control fat diet, but not in 60% fat-fed group. Handling of blood glucose during IPGTT was better in male *Nucb1* KO mice fed both diets, while such an effect was not observed in female KO mice. A significant two-way interaction of mice group and time on food and water intake value in 24 h was observed for male *Nucb1* KO mice fed 10% fat diet. The total physical activity during the dark phase and energy expenditure during the light phase showed a sex-specific pattern in male and female *Nucb1* KO mice fed 10% fat diet. Energy expenditure showed a sex-specific pattern in *Nucb1* KO mice during the dark phase. Moreover, adiposity increased in male *Nucb1* KO mice fed a high fat diet.

**Conclusions:** Our results indicate that the disruption of *Nucb1* leads to metabolic changes *in vivo*. The phenotype appears to depend on sex, age, diet, and the light-dark cycle. In conclusion, these outcomes furnish important evidence supporting critical roles for endogenous NUCB1 in energy homeostasis.

## 1. Introduction

Nucleobindin-1 (NUCB1) is a calcium and DNA binding protein that was originally reported in 1990. The peptide has both DNA and calcium-binding EF-hands [1, 2]. NUCB1 is a secreted peptide [3, 4], which is detected in many tissues in mammals, especially in humans and rodents [5, 6]. This peptide has been implicated in several biological processes, including calcium homeostasis [7], regulation of DNA [8], neuronal biology [9], hormone secretion, and protein folding [10]. These functions clearly place NUCB1 as an important mediator of cellular and physiological processes in mammals. Years later, another peptide was named nucleobindin 2 (NUCB2) due to its structural similarity with NUCB1 [11]. NUCB2 has calcium and DNA binding sites and is a secreted protein [1]. More recently, Oh-I and colleagues identified a bioactive peptide within NUCB2 that causes satiety and fat loss in rats and named it nesfatin-1 (NESF-1) [12]. Through immunoprecipitation and mass spectrometry, our lab discovered that a NESF-1-like peptide (NLP) is processed from NUCB1 [13]. Further research found that NLP is an anorexigenic peptide in rats [14] and goldfish [11] and has insulinotropic actions in mice [10]. We previously reported that NLP suppressed the basal and thyrotropin-releasing hormone (TRH)-induced prolactin expression in rat GH3 somatolactotrophs [15]. Moreover, NLP reduced growth hormone (GH) synthesis and secretion from rat somatotrophs while stimulated the synthesis of proopiomelanocortin (POMC) in mouse corticotrophs. This direct action of NLP on GH and POMC was mediated via the cAMP response element-binding protein (CREB) pathway [16]. Moreover, fluorescent-tagged NLP was able to bind to the surface of rat somatotrophs. These suggest a G coupled protein receptor (GPCR) mediated action of NLP in modulating GH [16]. While these evidence from multiple species and cells studies provide clear indications that NLP is a bioactive peptide that modulates whole-body energy homeostasis and hormone secretion, whether endogenous NUCB1 is critical for homeostasis is unknown. Therefore, the main goal of research outlined in this research was to address this gap in knowledge. We hypothesized that the disruption of endogenous *Nucb1* will negatively affect whole-body energy homeostasis in mice. Both male and female mice were studied separately. Our results indicate that the disruption of *Nucb1* results in mild metabolic defects in mice, and these effects occur in a sexually dimorphic manner.

## 2. Materials and Methods

### 2.1. Animals

Breeding pairs of homozygous C57BL/6NCrl-*NUCB1^em1(IMPC)Mbp^*/Mmucd, 043641-UCD mice from the University of California, San Diego (generated by Dr. Kent Lloyd, Mouse Biology Program, University of California-Davis, USA) were used to establish a colony of *Nucb1* disrupted mice. To generate these global *Nucb1* disrupted mice, the exon 4 and flanking splicing regions of *Nucb1* were constitutively deleted using CRISPR Cas9 gene editing technology in C57BL/6J mouse zygotes. All the animals were housed under 12 h light:12 h dark cycle (7 AM - 7 PM), humidity (30 – 60 %), and temperature (18 – 22 °C) controlled vivarium located in the College of Medicine Lab Animal Services Unit, University of Saskatchewan. All animal use and care protocols were prepared based on the guidelines of the Canadian Council for Animal Care and ARRIVE guidelines and they were approved by the University of Saskatchewan Animal Care Committee (Animal Care Protocol # 2012-0033).

### 2.2. Genotyping

Wildtype (WT) and knockout (KO) littermate genotypes were confirmed RT-PCR. Briefly, tissue samples were collected via tail clipping and DNA was extracted using Direct PCR Lysis Reagent (Cat no. 101 101 T, Viagen Biotech, USA), and 1.5 µL of the lysate from each sample was used for PCR. The Taq DNA Polymerase recombinant PCR kit (Cat no. 10342-020, Thermo Fisher Scientific, USA) was used following the manufacturer’s protocol. For the *Nucb1* gene, pairs of forward (GTTGTCTCTGTTGAGTATTGAAAGACAGG) and reverse primers (GGATACGAGCTCAGGTCACCAGG) were designed by MMRRC at University of California (Davis) to specifically amplify either the WT or the KO allele in separate PCR amplifications. The PCR products were separated on 1.5% agarose gel, and the images were captured using the ChemiDoc MP Imaging System (Bio-Rad, Canada).

After confirmation of the homozygous KO genotype, age-matched WT and *Nucb1* KO mice for experiments were moved to the Animal Care Unit of the Western College of Veterinary Medicine (University of Saskatchewan) and maintained under the same conditions. These mice were used for two different studies, explained below. After finishing all experiments, pancreas (whole tissue) and large intestine (transverse colon) samples were collected from each mouse after euthanasia to re-confirm the genotype of each animal. In this case, total RNA was extracted, and cDNA was made as mentioned below. Finally, 1 µL of cDNA of each sample was subjected to PCR amplification, PCR product separation in agarose gel and imaging, as explained above.

### 2.3. Diets

The first set of studies were carried out on age-matched mice fed a standard rodent chow diet (Cat no. 500I, LabDiet, 7.94% carbohydrate, 28.67% protein, 13.38% fat, Energy density = 4.09 Kcal/g). Second set of studies were conducted on age-matched mice fed a control fat diet (Cat no. D12450B, Research Diets Inc., 70% carbohydrate, 20% protein, 10% fat, Energy density = 3.82 Kcal/g) and high-fat diet (Cat no. D12492, Research Diets Inc., 20% carbohydrate, 20% protein, 60% fat, Energy density = 5.21 Kcal/g). The full list of ingredients for 10% and 60% fat diets are provided in ***Supplementary Table 1***. Bodyweight, food intake and blood glucose of all mice were weekly monitored as explained below. In addition, whole-body metabolic parameters were measured using the Comprehensive Lab Animal Monitoring System (CLAMS, Columbus Instruments, USA), as well as the ability to handle glucose was tested (see below). After experiments were done, mice were euthanized by cervical dislocation. Bodyweight of mice was noted, and liver and weight adipose tissue were weighed and collected.

### 2.4. Food intake, blood glucose, bodyweight – weekly monitoring

To determine the phenotypic changes in male and female *Nucb1* KO mice versus WT littermates fed either standard chow diet or high-fat diet, blood glucose level was recorded weekly after fasting mice for 4-h using glucometer for 24-25 weeks (OneTouch Ultra^®^, LifeScan, Canada). In addition, food intake and bodyweight gain of mice were also monitored weekly using a digital scale. Mice were age-matched, and the monitoring started around 10 weeks post-weaning for chow-fed mice and 13 weeks post-weaning for high-fat-fed mice.

### 2.5. Oral or intraperitoneal glucose tolerance tests (O/IPGTT)

To elucidate whether genetic disruption of *Nucb1* affect orally administered or intraperitoneally injected glucose tolerance, OGTT and IPGTT were performed, respectively. These tests were performed (littermate controls and *Nucb1* KO mice, n ≥ 6) in age-matched mice (33-34 weeks old for chow-fed mice and 25-26 weeks old for high-fat-fed mice). All animals were fasted 4 h (9 AM) before the commencement of oral gavage (OGTT) or IP administration (IPGTT) at 1 PM. For OGTT, 2 g/kg bodyweight of D-Glucose (Fisher Chemical, #D16-500) dissolved in sterile water (Gavage volume: 200 µL) was administered to mice using insulin syringes (BD, #324911) attached to animal gavage needles (Popper and Sons, USA, #9921). For IPGTT, 2 g/kg bodyweight of D-Glucose dissolved in sterile saline (injection volume: 200 µL) was administered to mice using insulin syringes attached to a 27G needle in the lower right quadrant of the abdomen to avoid damage to the urinary bladder, caecum, and other abdominal organs. For both tests, blood glucose via tail snip was measured at time points 0, 10, 20, 30-, 60-, 90- and 120-min post-administration using a glucometer. Each bleeding took 30-60 s, and mice were returned to their respective cages until the next bleed. The area under the curve (AUC) was calculated using GraphPad Prism Version 8.0 (GraphPad Inc., USA) [The formula (concentration of glucose at T1 + concentration of glucose at T2)/2·(T2-T1), where T1 and T2 are two consecutive time points].

### 2.6. Assessment of whole-body energy homeostasis using the comprehensive lab animal monitoring system (CLAMS)

The whole-body metabolic parameters were measured in both male and female littermate control and *Nucb1* KO mice using the CLAMS (Columbus Instruments, USA). Mice were transferred to CLAMS cages to acclimate for 3 days before monitoring feeding, drinking, and other metabolic parameters over 24 h (one dark phase and one light phase). CLAMS monitoring was completed twice each on chow-fed mice (20-21 weeks old) and high-fat-fed mice (28-29 weeks old), and then data from each diet group repetition were pooled from two studies (n = 4 mice/group maximum tested per study) for each sex. The CLAMS system was calibrated according to the manufacturer’s instructions prior to beginning the experiments. This system is equipped with an open-circuit calorimeter for measuring cumulative oxygen (O_2_), O_2_ consumption (VO_2_), carbon dioxide production (VCO_2_), and energy expenditure (Heat). The respiratory exchange ratio (RER) was calculated by dividing VCO_2_ by VO_2,_ with values ranging from 0.7 to 1.0. A shift in RER value towards 0.7 indicates that fat is the major substrate used for energy production, while a shift towards 1.0 indicates that carbohydrate is the main substrate used. The percent of fat and carbohydrates utilized for energy by each animal was determined from the RER. The energy derived from carbohydrates or fats was extrapolated from RER data by employing the methods of Mclean and Tobin [17], using the equation and methods described in the protocols provided by the Columbus instruments (http://www.colinst.com). Accordingly, the equations are: energy derived from carbohydrate = Percentage of carbohydrate/100 × Heat. Percentage carbohydrate values are based on the RER values (RER range; % carbohydrate: 0 – 0.7285: 0; 0.7285 – 0.775: 14.7; 0.775 – 0.825: 31.7; 0.825 – 0.875: 48.8; 0.875 – 0.925: 65.9; 0.925 – 0.975: 82.9; Above 0.975: 100). Percentage fat = 100 - % carbohydrate. Energy derived from fat = Percentage of fat / 100 × Heat. Each cage is also equipped with a system of infrared (IR) beams that detect animal movement in the three planes/axes (X, Y and Z). We measured the total physical activity as the sum of number of activities recorded as X+Y+Z axes IR beam breaks. To adhere to the figure limits of this journal, the most significant results from the CLAMS measurements are presented.

### 2.7. RNA extraction, cDNA synthesis and qPCR

Different tissues from chow-fed mice were collected, and total RNA was extracted, cDNA was made, and qPCR was performed according to the published methods [18]. Briefly, tissues were homogenized in 1 mL of RiboZol (Cat no. N580, VWR, USA), and then total RNA was precipitated in 500 μL of isopropanol. After the final washing step with 75% ethanol twice, RNA pellets were suspended in RNase-free water. The concentration of total RNA was measured using the NanoDrop2000 (Thermo Fisher Scientific, USA), and RNA purity was assessed by 260/280 and 260/230 ratios. Total RNA (1 μg) was reverse transcribed to cDNA using an iScript cDNA synthesis kit (Cat no. 170884, BioRad, USA), following the manufacturer’s instructions. qPCR was conducted using the CFX Connect Optic module (BioRad, USA) using SensiFAST SYBR No-ROX Kit (Cat no. BIO-98050, Bioline, Froggabio, Canada). The amplification reaction contained 1 μL of diluted cDNA, 0.5 μL of primer mix (forward and reverse) at 10 μM, 5 μL of SYBR green PCR master mix and water up to the total volume of 10 μL. The amplification efficiency for each primer set was obtained from a serial dilution of cDNAs with a dilution factor of 4. Those primers with reaction efficiency of 90–110%, with a single disassociation peak of PCR products and the absence of primer-dimer were considered in this study. The dilution factor of cDNA samples was selected based on the best threshold cycle (CT) range for each primer set. The primer sequences and their annealing temperatures are listed in ***Supplementary Table 2***. The thermal cycling set-up for all genes was the following: denaturation (95 ^◦^C for 5 s), annealing (specific for each primer for 25 s) and elongation (60 ^◦^C for 25 s), 35 cycles. Three negative controls, including no template DNA (NTC control), no reverse transcriptase control from cDNA synthesis process (RTC control), and a nuclease-free water sample (PCR control) were also included for each gene. The transcript abundance was calculated based on the Pfaffl method based on gene-specific efficiencies relative to the geometric means of the two housekeeping genes.

### 2.8. Statistical analysis

Two-way mixed ANOVA (mice strain [group] and time factor) was used to compare the mean differences between WT and *Nucb1* KO mice on the weekly measured bodyweight, blood glucose, and food intake during 24-25 weeks, as well as to compare the food and water intake during 24 h in CLAMS studies. Before running the two-way mixed ANOVA, the normal distribution of the data and the presence of outliers were evaluated using Shapiro-Wilk test and ROUT methods, respectively. Sphericity was not assumed following the recommendation of Maxwell and Delaney [19], and the Geisser and Greenhouse epsilon hat method was used to adjust the results considering the epsilon value. Bonferroni’s multiple comparison test was performed as a *post hoc* analysis to compare the dependent variable results between mice strain (group) in each time-point. Unpaired samples *t*-test was used for analyzing the differences in energy expenditure, energy derived from carbohydrates or fat and total physical activity in CLAMS studies, as well as to compare AUC in the glucose tolerance tests (IPGTT and OGTT). All analysis were carried out using GraphPad Prism Version 8.0 (GraphPad Inc., USA) and statistical differences were considered at *p* < 0.05.

Chow fed and high-fat-fed mice data were also analyzed using generalized estimating equation (GEE) statistical method. In this test, mouse ID was considered as subject effect. The time blocks/duration varied based on the nature of experiment (week/mice age for weekly monitoring of bodyweight, food intake and blood glucose, hour for CLAMS as well as time point for glucose tolerance tests) that was considered as within-subjects factor and as a covariate in predictor section of GEE commands. The majority of predictors were binary variables, including female and male for mice sex, WT and *Nucb1* KO for mice group (presence or absence of an undisrupted *Nucb1* gene), light and dark phases for day phase. As mentioned above, diet is a binary factor for high fat diet experiments, which have two levels, control fat diet (10% fat) and experimental fat diet (60% fat). The first order of autoregressive model (AR1) was chosen as an ideal working correlation matrix for all experiments. Repeated measures were assumed to have AR1 relationship, which means the current values are a function of prior values plus error. This assumes time series data with equal intervals. This model is a common choice in longitudinal models such as time series experiments. The scale response was chosen as linear. First, the general effect of predictors on dependent variables were examined and then two-, three- and four-way interactions of predictors in the model were checked, respectively.

## 3. Results

### 3.1. Genotyping

PCR on genomic DNA provided bands of the expected size (397 bp) and confirmed the disruption of *Nucb1* in homozygous mice, compared to WT littermates (611 bp) (**Figure 1A**). Some samples from *Nucb2* KO mice were used to validate the results. Since the genetic disruption in *Nucb2* was not within the location where primers are targeted, we expected to observe the same band size as WT for this group.

**Figure 1.**
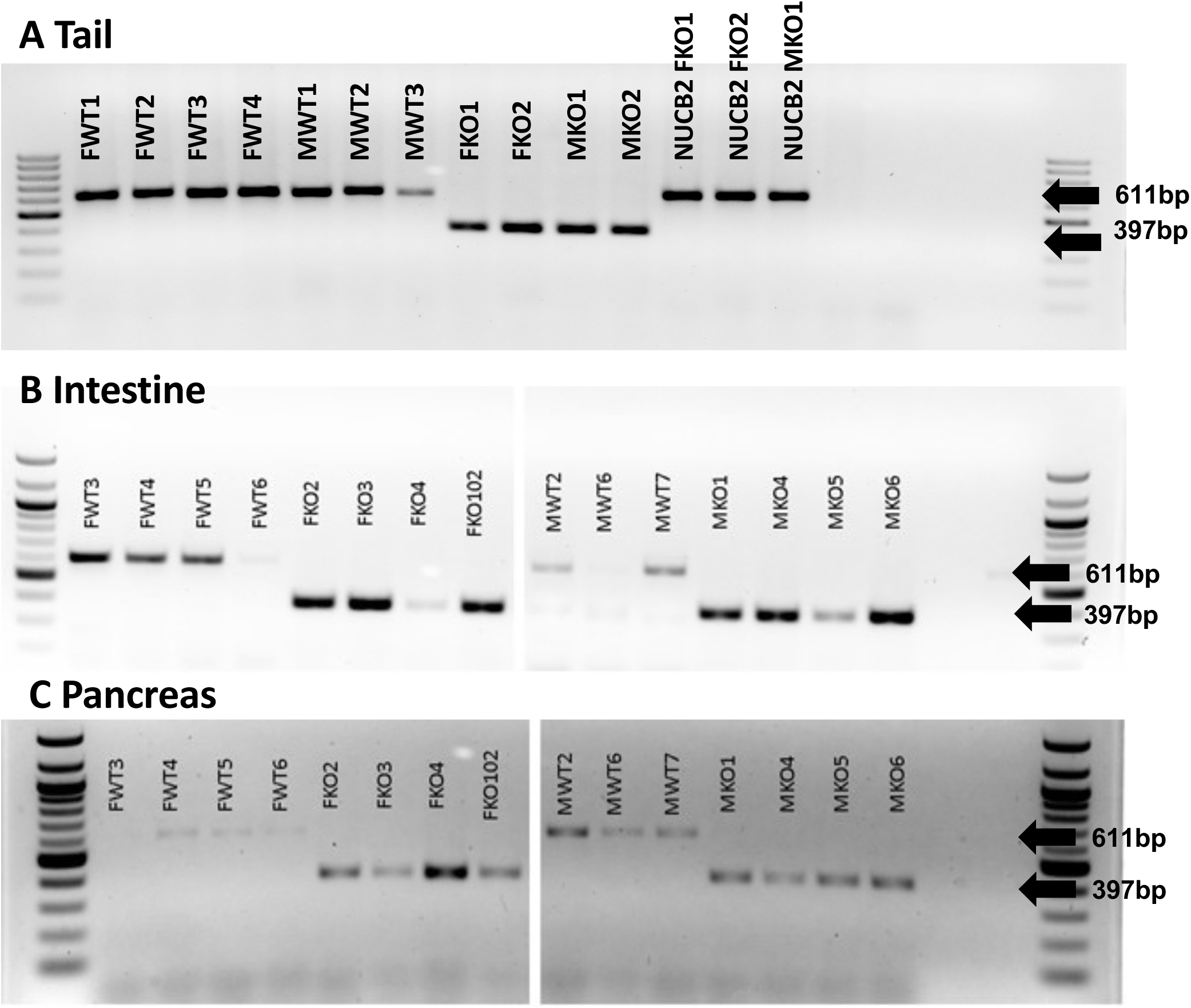
RT-PCR analysis of tail (A), intestine (B) and pancreas (C) to confirm the disruption of *Nucb1* gene in *Nucb1* KO mice and to compare it with WT mice. The band corresponding to 397 bp shows the disruption of *Nucb1* in mice and band corresponding to 611 bp shows intact *Nucb1* gene in WT mice.

The analysis of the intestine and pancreas showed the predicted band corresponding to the 397 bp in *Nucb1* mice. Although the band around 611 bp was not detected in any male or female WT mice, the band corresponding to 397 bp was absent in all WT mice (**Figure 1B** and **Figure 1C**).

### 3.2. Weekly bodyweight, food intake, and basal blood glucose monitoring

#### Weekly bodyweight

Although the bodyweight of chow fed male *Nucb1* KO mice was initially lower than that of WT littermates in the first monitoring phase (weeks 11-19), from week 20 the bodyweight has normalized, and no significant differences were observed throughout the rest of the experiment (**Figure 2A**). In contrast, the disruption of *Nucb1* in female mice did not affect bodyweight throughout the experiment (**Figure 3A**). No significant differences in bodyweight were found in male and female *Nucb1* KO mice fed either 10% or 60% fat diet during the monitoring (**Figure 4A** and **Figure 5A**).

**Figure 2.**
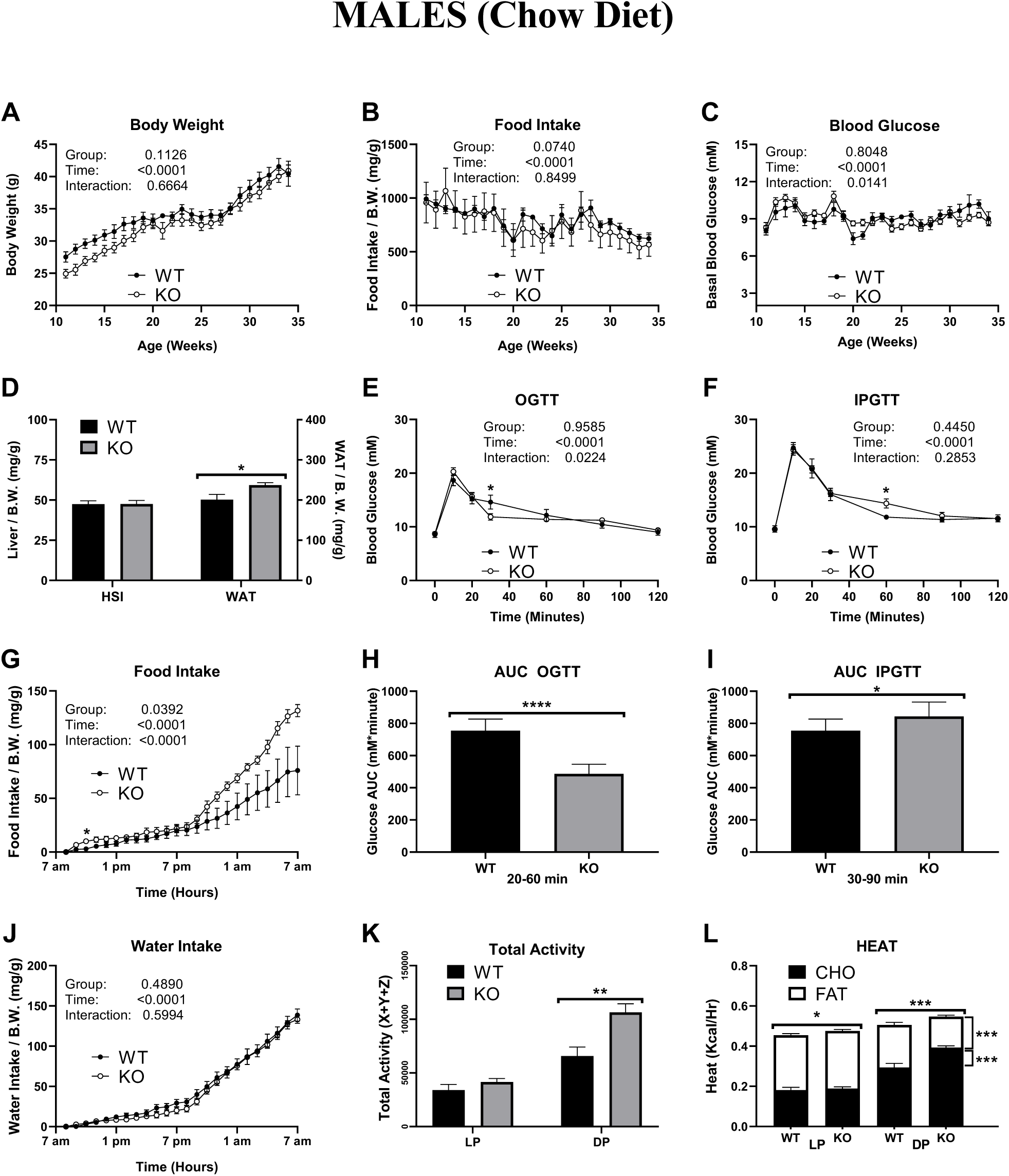
General characterization of chow-fed male WT and *Nucb1* KO mice. Normal distribution of the data, the homogeneity of variances and covariances were determined using Shapiro-Wilk test, Levene’s test, and Box’s test, respectively. The two-way mixed ANOVA was performed. A Bonferroni’s multiple comparison test was also performed as a post hoc analysis to compare the dependent variable results between mice strain (group) in each time point. Unpaired t-test was used for analyzing the differences in energy expenditure, energy derived from carbohydrates or fat and total physical activity in CLAMS studies, as well as to compare AUC in the glucose tolerance tests (IPGTT and OGTT). All the analyses were carried out using GraphPad Prism 8.0, and statistical differences were considered at p < 0.05. The number of mice in each of the experiments is provided below: Weekly tracking: WT n = 7; KO n = 12, CLAMS: n = 6-8 mice/group/sex (20-21 weeks old), Glucose tolerance tests (GTTs): WT n = 7; KO n = 10-12

**Figure 3.**
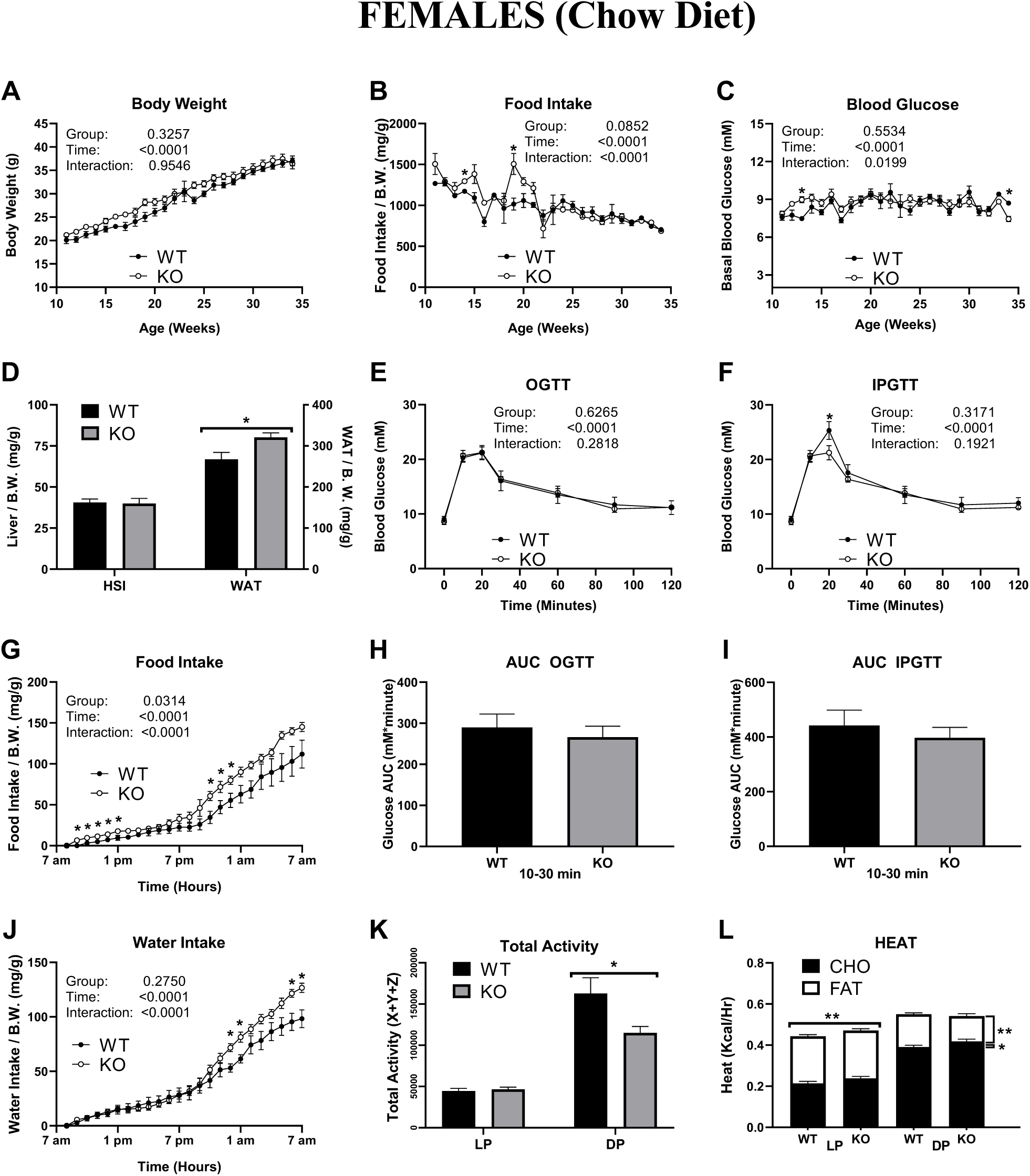
General characterization of chow-fed female WT and *Nucb1* KO mice. Normal distribution of the data, the homogeneity of variances and covariances were determined using Shapiro-Wilk test, Levene’s test, and Box’s test, respectively. The two-way mixed ANOVA was performed. A Bonferroni’s multiple comparison test was also performed as a post hoc analysis to compare the dependent variable results between mice strain (group) in each time point. Unpaired t-test was used for analyzing the differences in energy expenditure, energy derived from carbohydrates or fat and total physical activity in CLAMS studies, as well as to compare AUC in the glucose tolerance tests (IPGTT and OGTT). All the analyses were carried out using GraphPad Prism 8.0, and statistical differences were considered at p < 0.05. The number of mice in each of the experiments is provided below: Weekly tracking: WT n = 7-8; KO n = 8-12, n was varied due to variation in sample size for some weeks, CLAMS: n = 6-8 mice/group/sex (20-21 weeks old), Glucose tolerance tests (GTTs): WT n = 6; KO n = 7

**Figure 4.**
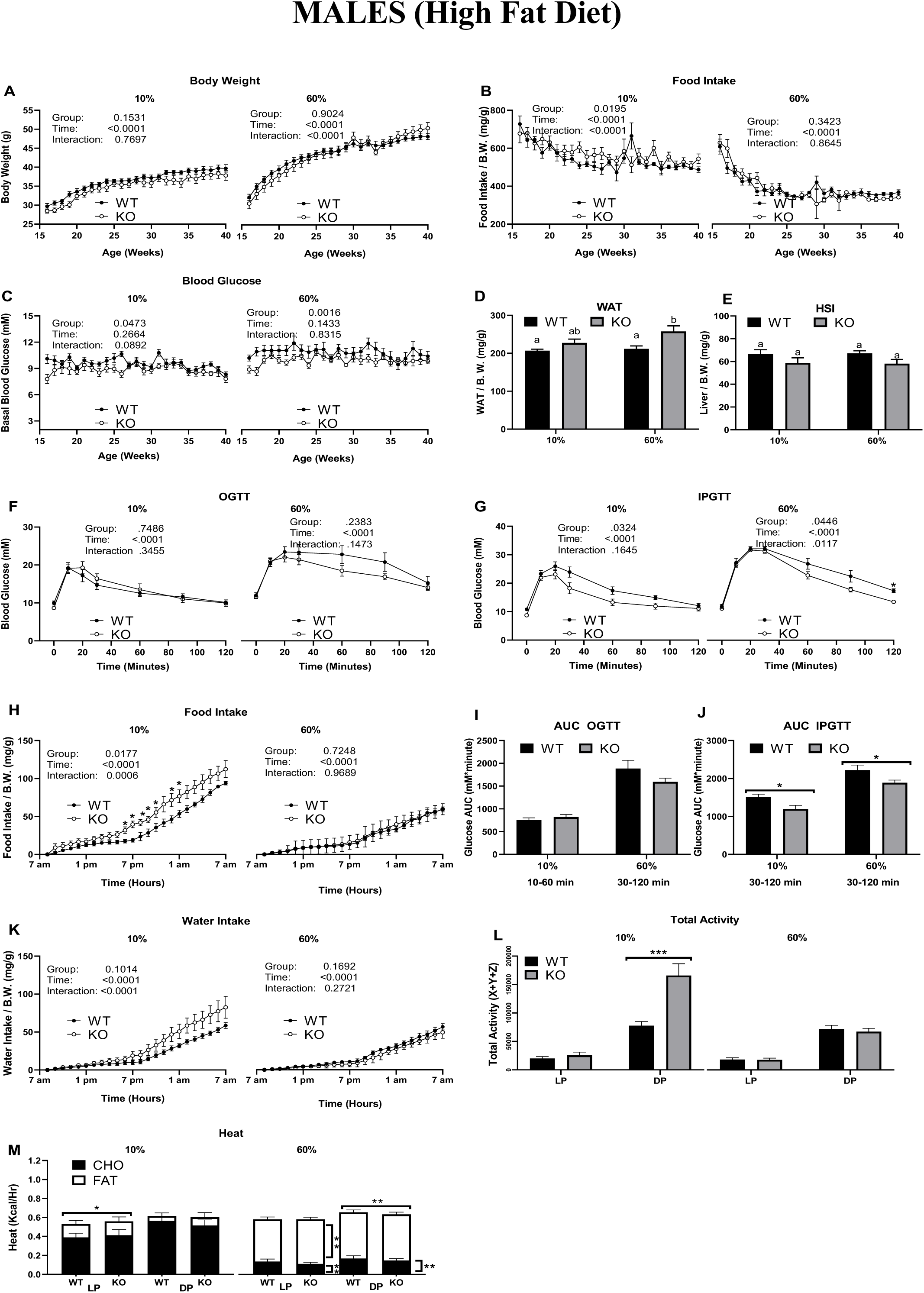
General characterization of control and high-fat-fed male WT and *Nucb1* KO mice. Normal distribution of the data, the homogeneity of variances and covariances were determined using Shapiro-Wilk test, Levene’s test, and Box’s test, respectively. The two-way mixed ANOVA was performed. A Bonferroni’s multiple comparison test was also performed as a post hoc analysis to compare the dependent variable results between mice strain (group) in each time point. Unpaired t-test was used for analyzing the differences in energy expenditure, energy derived from carbohydrates or fat and total physical activity in CLAMS studies, as well as to compare AUC in the glucose tolerance tests (IPGTT and OGTT). All the analyses were carried out using GraphPad Prism 8.0, and statistical differences were considered at p < 0.05. The number of mice in each of the experiments is provided below: Weekly tracking: 10% fat fed mice WT n = 10-12; KO n = 10-14, 60% fat fed mice WT n = 9-11; KO n = 10-13n was varied due to variation in sample size for some weeks, CLAMS10% fat fed mice WT n = 6-8; KO n = 6-9, 60% fat fed mice WT n = 6-7; KO n = 8-9 mice/group/sex (28-29 weeks old), Glucose tolerance tests (GTTs): 10% fat fed mice WT n =12; KO n = 10-11, 60% fat fed mice WT n = 8; KO n = 13

**Figure 5.**
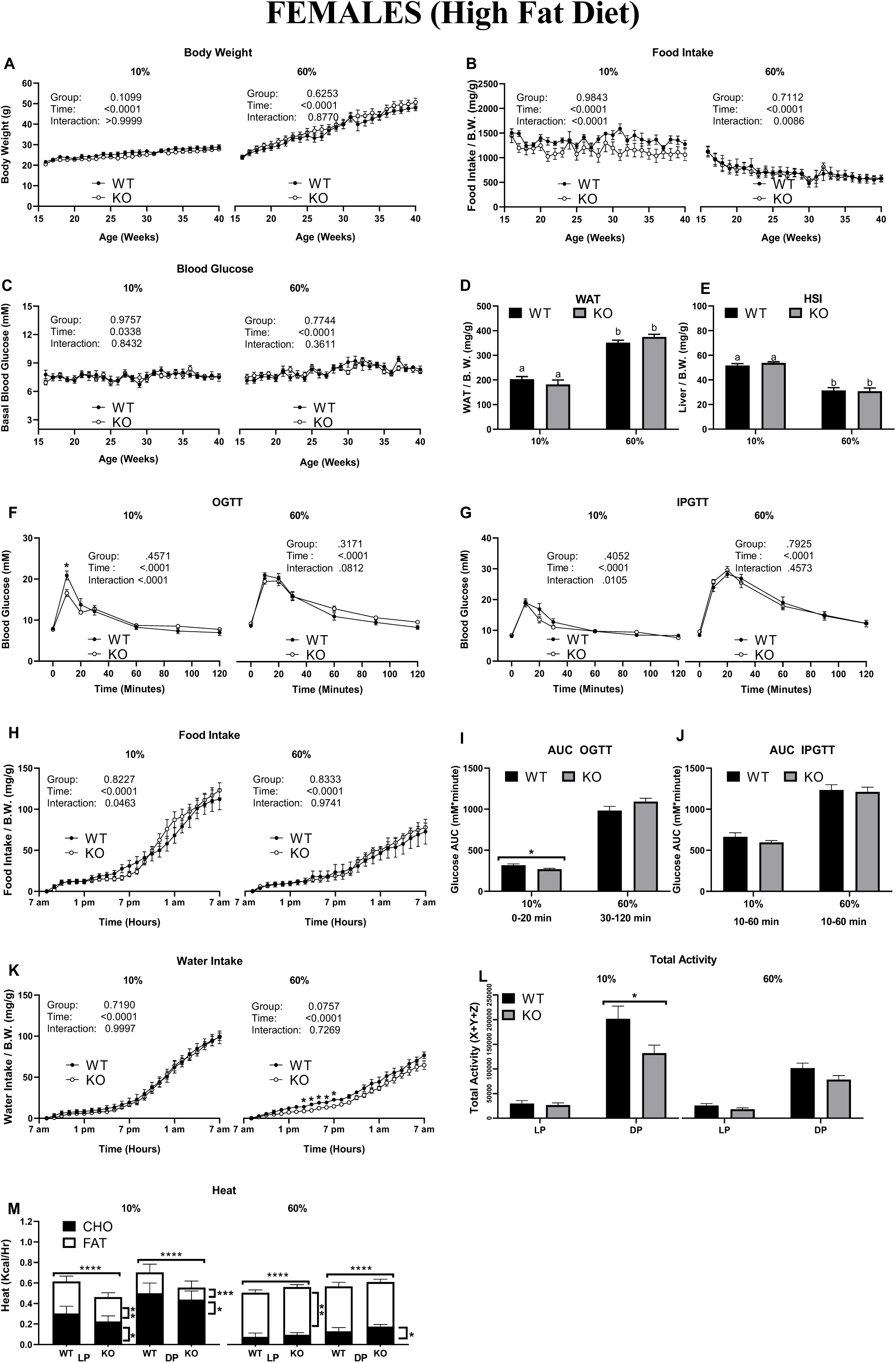
General characterization of control and high-fat-fed female WT and *Nucb1* KO mice. Normal distribution of the data, the homogeneity of variances and covariances were determined using Shapiro-Wilk test, Levene’s test, and Box’s test, respectively. The two-way mixed ANOVA was performed. A Bonferroni’s multiple comparison test was also performed as a post hoc analysis to compare the dependent variable results between mice strain (group) in each time point. Unpaired t-test was used for analyzing the differences in energy expenditure, energy derived from carbohydrates or fat and total physical activity in CLAMS studies, as well as to compare AUC in the glucose tolerance tests (IPGTT and OGTT). All the analyses were carried out using GraphPad Prism 8.0, and statistical differences were considered at p < 0.05. The number of mice in each of the experiments is provided below: Weekly tracking: 10% fat fed mice WT n = 9-10; KO n = 11-15, 60% fat fed mice WT n = 10-11; KO n = 12-14 n was varied due to variation in sample size for some weeks, CLAMS10% fat fed mice WT n = 6-8; KO n = 7-9, 60% fat fed mice WT n = 6-7; KO n = 7-8 mice/group/sex (28-29 weeks old), Glucose tolerance tests (GTTs): 10% fat fed mice WT n = 10; KO n = 14, 60% fat fed mice WT n = 11; KO n = 14

#### Weekly food intake

The genetic disruption of *Nucb1* in male mice did not alter food intake during the 6-month monitoring when compared to WT controls (**Figure 2B**). Female *Nucb1* KO mice showed higher food intake at some weeks (**Figure 3B**). Hence, a significant interaction between female mice groups and time was observed for food intake (p <0.0001). The male *Nucb1* KO mice fed control and high-fat diet did not reveal higher food intake during weekly monitoring compared to their corresponding WT littermates (**Figure 4B**). Interestingly, a significant interaction between 10% fat-fed male mice and time on food intake was observed (p <0.0001). Although the disruption of *Nucb1* in female mice fed control and high-fat diets did not alter food intake during weekly monitoring (**Figure 5B**), a significant interaction between time and female mice groups on food intake was observed for both control and high-fat-fed mice.

#### Weekly basal blood glucose levels

The genetic disruption of *Nucb1* in male mice did not alter the blood glucose level (**Figure 2C**). However, a significant interaction between time and male mice groups on blood glucose levels was found (p = 0.0141). In contrast, in female mice, significant differences were found in the blood glucose level at one specific point (i.e., week 13), being higher in KO mice compared to the WT littermates and for week 34, blood glucose is higher in WT compared to the KO mice (**Figure 3C**). A significant interaction effect between female mice groups and time on blood glucose level was found (p = 0.0199). As shown in **Figure 4C**, the general group effect on basal blood glucose levels in male mice was significant for both control and high-fat diet groups. Whereas, the general group effect on basal blood glucose levels in female mice fed control and high-fat diet was not significant which shows that the effect on blood glucose levels might be sexually dimorphic in *Nucb1* disrupted mice. No changes were observed for basal blood glucose level of 60% fat-fed KO mice (**Figure 5C**). Comprehensive GEE analysis showed when all predictors (sex, age, diet, and mice groups) are gathered in the united data matrix, the general effects of *Nucb1* genetic disruption (mice group) affected basal blood glucose significantly.

The results of GEE analysis of chow-fed mice and high fat-fed data are summarized in ***Supplementary Table 3*** and ***Table 4***, respectively. As shown in ***Table 3***, the interaction of mice sex, group, and age on weekly monitoring of chow-fed mice parameters was significant, indicating the effect of genetic disruption of *Nucb1* on bodyweight, food intake and basal blood glucose level of chow-fed mice is sex- and age-dependant. Moreover, the GEE analysis showed significant general effects of sex, age, diet and mice group and various interaction combinations of these predictors on weekly monitoring parameters in control and high-fat-fed mice (***Table 4***). These results suggest sex-, age-, mice strain- and diet-specific effects when *Nucb1* is disrupted.

### 3.3. Glucose tolerance tests (OGTT and IPGTT)

#### OGTT

We determined whether the genetic disruption of *Nucb1* affects oral glucose handling in mice. Blood glucose level was significantly lower at 30 min in male *Nucb1* KO mice after the oral administration of 2g glucose/kg bodyweight compared to the littermate control group (**Figure 2E**). Consistent with this result, the area under the curve (AUC) analysis shows that male *Nucb1* KO mice handled blood glucose levels better during 20-60 min of the OGTT (**Figure 2H**). However, this effect occurred in a sex-specific manner because no such effects were found in female *Nucb1* KO mice, whose handling of blood glucose was not different from littermate controls (**Figure 3E** and **Figure 3H**). The disruption of *Nucb1* did not affect blood glucose handling in high-fat-fed mice of either sex after the oral administration of 2g glucose/kg bodyweight (**Figure 4F and Figure 5F**). However, a reduced blood glucose level in female *Nucb1* KO mice fed control fat diet at 10 min of OGTT was observed. The measurement of AUC for 10% fat-fed female mice illustrates the same result during 0-20 min of OGTT (**Figure 5F** and **Figure 5I**). In addition, a significant interaction between female mice group and time point was also observed (p<0.0001) (**Figure 5F**). In contrast, male *Nucb1* KO mice did not show significant changes in blood glucose level when compared to WT littermates (**Figure 2C**).

#### IPGTT

Male *Nucb1* KO mice had higher blood glucose level at 60 min after the IP administration of glucose compared to WT littermate. Consistently, AUC measurement of showed higher value during 30-90 min of IPGTT (**Figure 2F** and **Figure 2I**). In contrast, the genetic disruption of *Nucb1* caused lower blood glucose levels in female mice at 20 min of IPGTT (**Figure 3F**). However, the measurement of AUC illustrates the no significant difference between female WT and KO during 10-30 min of IPGTT (**Figure 3I**). Moreover, male *Nucb1* KO mice fed control or high-fat diets had lower blood glucose compared to WT during 30-120 min of IPGTT compared to WT fed the same diets (**Figure 4J**). However, this difference in blood glucose was only significant for male *Nucb1* KO mice at 120 min of IPGTT (**Figure 4G**). A significant general effect of male mice group for both diets was detected. Besides, a significant interaction of mice group and time point on blood glucose level was observed for male mice fed high-fat diet (**Figure 4G**). In contrast, female *Nucb1* KO mice fed control and high-fat diet did not reveal any significant changes in blood glucose handling during IPGTT. A significant interaction (p=0.0105) was observed for 10% fat-fed female mice (**Figure 5G**).

Overall, comprehensive GEE analysis on OGTT results of male and female mice fed a chow diet showed that the glucose handling in the presence of *Nucb1* gene disruption occurs in a sex- and time point-specific manner. Oral glucose handling in control and high-fat diet groups is influenced by sex, fat percentage, time point and presence/absence of *Nucb1* disruption.

The general effect of time point and interaction of time point, sex, and mice group on blood glucose handling in chow-fed mice got significant during IPGTT. The general effect of *Nucb1* gene disruption (group), time point, sex, and fat percentage on blood glucose handling of mice fed control and high-fat diet was detected as significant. Moreover, the different interactions of sex, group, time point and diet in this group got significant. Although the comprehensive GEE analysis did not demonstrate a significant general effect of *Nucb1* disruption on glucose handling during OGTT and IPGTT except for high-fat-fed mice during IPGTT, it showed that the genetic disruption of *Nucb1* affects glucose handling in a sex-, diet- and post-glucose administration time-dependent manner (***Supplementary Tables 3 - 4***).

### 3.4. Assessment of whole-body energy homeostasis using comprehensive lab animal monitoring system (CLAMS)

#### Twenty-four-hour food intake

In general, the disruption of *Nucb1* in male and female mice altered cumulative food intake, which was found higher in KO mice when compared to littermate WT mice (p = 0.0392 and 0.0314, respectively) (**Figure 2G** and **Figure 3G**). Although this difference for male mice is more evident during the dark phase (7 PM - 7 AM), a significant difference in food intake of male mice was only found at the beginning of the light period (9 AM). In contrast, significant differences in food intake of female mice were found between groups at different time points in both light and dark phases. Due to time-specific responses, a significant interaction between mice groups and time was observed on cumulative food intake in both sexes (p < 0.0001).

The genetic disruption of *Nucb1* altered food intake in males fed a control fat diet (p=0.0177). A significant interaction on food intake was observed for these male mice with higher food intake during the dark phase (**Figure 4H**). In contrast, no difference in food intake was observed between male *Nucb1* KO and WT mice fed a high-fat diet during the light or dark phases (p=0.7248, **Figure 4H**). Food intake of female *Nucb1* KO mice fed both fat diets was similar to that of female WT mice. A significant interaction of mice group and time on food intake was only observed in female mice fed control (fat) diet (p=0.0463, **Figure 5H**).

#### Twenty-four-hour water intake

Cumulative water intake was significantly different among female groups at different time points of the dark phase (**Figure 3J**), being higher for *Nucb1* KO mice. However, no such differences were observed during the light period, and no general group effect was detected (p = 0.2750), but a significant interaction between group and time was found (p < 0.0001). The effect on water intake due to *Nucb1* disruption was sexually dimorphic due to *Nucb1* disruption because no significant differences were observed in cumulative water intake between groups in male mice (**Figure 2J**). Furthermore, water intake consumption in *Nucb1* KO mice were similar to that of WT mice of both sexes fed a control fat diet during light and dark phases. The cumulative water intake in female WT mice fed a high-fat diet was significantly higher during limited hours in the late light phase. However, no changes were found in male mice fed the same diet. A significant interaction between male mice groups fed the control fat diet and time on water intake was detected (p<0.0001), whereas such an interaction was not observed in female mice fed the same diet (**Figure 4K** and **Figure 5K**).

#### Twenty-four-hour physical activity

The disruption of *Nucb1* induced a significant increase in physical activity in male mice during the dark phase compared to WT littermates (**Figure 2K**). In contrast, physical activity in female mice displayed the opposite (decrease) of what was found with male mice in the dark phase (**Figure 3K**). Thus, here also, a sexually dimorphic effect was found due to the genetic disruption of *Nucb1*. Moreover, male, and female *Nucb1* KO mice fed control fat diet displayed similar results on physical activity (increase in male *Nucb1* KO and decrease in female *Nucb1* KO mice in dark phase) of what was found in chow-fed mice groups (**Figure 4L** and **Figure 5L**). These findings provide evidence on the sex-specific role of *Nucb1* disruption for physical activity when the fat percentage in diet is low. In contrast, high-fat-fed *Nucb1* KO mice did not show any significant physical activity compared to their WT littermates (**Figure 4L** and **Figure 5L**).

#### Twenty-four-hour energy expenditure

Our results indicate that male *Nucb1* KO mice fed chow diet had higher energy expenditure (heat) than WT littermates in both light and dark phases (**Figure 2L**) but an increase in energy expenditure in female *Nucb1* KO mice was found only during the light phase (**Figure 3L**). Energy expenditure showed an increasing pattern during the light phase in male *Nucb1* KO mice fed 10% fat while an opposite (decrease) effect in male KO groups fed high fat diet during the dark phase (**Figure 4M**). In female mice, energy expenditure showed a decreasing pattern in KO groups fed control fat diet in both phases, but an increasing pattern in those fed high fat diet in both phases (**Figure 5M**).

#### Twenty-four-hour energy derived from carbohydrates and fat

Interestingly, our results demonstrate that the catabolism of neither carbohydrates (CHO) nor fat was affected during the light phase between groups of either sex. The use of CHO in both male and female *Nucb1* KO mice was increased in the dark phase while fat catabolism was reduced compared to WT mice (**Figure 2L** and **Figure 3L**). The disruption of *Nucb1* decreased energy-derived from fat and CHO in female mice fed control fat diets in both phases (**Figure 5M**). High fat-fed male *Nucb1* KO mice had lower energy-derived from CHO during both phases, but female KO mice displayed an increase in energy derived from CHO in both phases. Moreover, energy-derived from fat was higher in female *Nucb1* KO mice in both phases, but in male KO mice was lower only in dark phase (**Figure 4M** and **Figure 5M**).

Furthermore, GEE analysis showed the general effects of sex and age on weekly monitoring of food intake and bodyweight in chow-fed mice were significant. The general effect of sex on the weekly monitoring of basal blood glucose levels of chow-fed mice was also significant. Moreover, the general effects of sex, age and diet on food intake and bodyweight were significant. The general effect of *Nucb1* gene disruption on the weekly monitoring of blood glucose of high-fat-fed mice were significant. Regarding the OGTT, the general effect of sex and time point in chow-fed mice and the general effect of sex, time point and diet on oral blood glucose handling were statistically significant. The general effects of all factors (sex, time group, *Nucb1* disruption, diet) on IP glucose handling were significant for high-fat-fed mice.

GEE analysis of chow-fed mice data showed that the genetic disruption of *Nucb1* affected cumulative food intake in mice, and the general effects of sex and/or dark/light phase for other CLAMS parameters, including water intake, energy expenditure and total physical activity in the mice fed the same diet were significant. Overall, the GEE analysis supports significant 3-way interactions between the mice group, light phase, and mice sex on micro measurement of food intake and water intake, energy expenditure and total physical activity in chow-fed mice. This suggests that the genetic disruption of *Nucb1* affected whole-body energy metabolism of chow-fed mice in a sex- and light phase-specific manner.

GEE analysis supports the results obtained from control and high-fat-fed mice, including significant interactions between mice group, day phase, sex, and diet on CLAMS parameters. This suggests that metabolic functions of obese mice were affected in *Nucb1* disrupted mice in a sex-, diurnal phase- and diet-specific manner.

### 3.5. Liver and adipose tissue weight

Our results showed that although HSI was not different between both groups (i.e., WT and *Nucb1* KO) in mice of either sex fed chow diet, WAT was higher in *Nucb1* KO mice in both sexes (**Figure 2D** and **Figure 3D**). Like chow-fed mice results, the HSI index was unaffected by the genetic disruption of *Nucb1* in control and high fat-fed male and female mice (**Figure 4E** and **Figure 5E**). Unlike increased adiposity seen in both sexes of chow-fed KO mice, adiposity was higher only in *Nucb1* disrupted male mice fed 60% fat diet (**Figure 4D**).

### 3.6. Metabolic gene expression in tissues of chow-fed mice

As shown in **Figure 6**, the abundance of *preproghrelin* and *Nucb2* mRNAs in the stomach of male *Nucb1* KO mice was increased, while no changes were observed in the female group (**Figure 6B**). In the duodenum, the *Gip* and *Glp-1* precursor mRNA levels were increased in female *Nucb1* KO mice without any change in male mice (**Figure 6L**). *Nucb2* abundance in the jejunum of *Nucb1* KO mice showed a sex-specific pattern with a decrease in male KO and an increase in female KO groups (**Figure 6G**). *Ucp-1* mRNA in brown fat was enhanced in both sexes of *Nucb1* KO mice (**Figure 6H**). *Glut1*, *Glut2* and *Nucb2* mRNAs were unaffected by the disruption of *Nucb1* in muscle samples.

**Figure 6.**
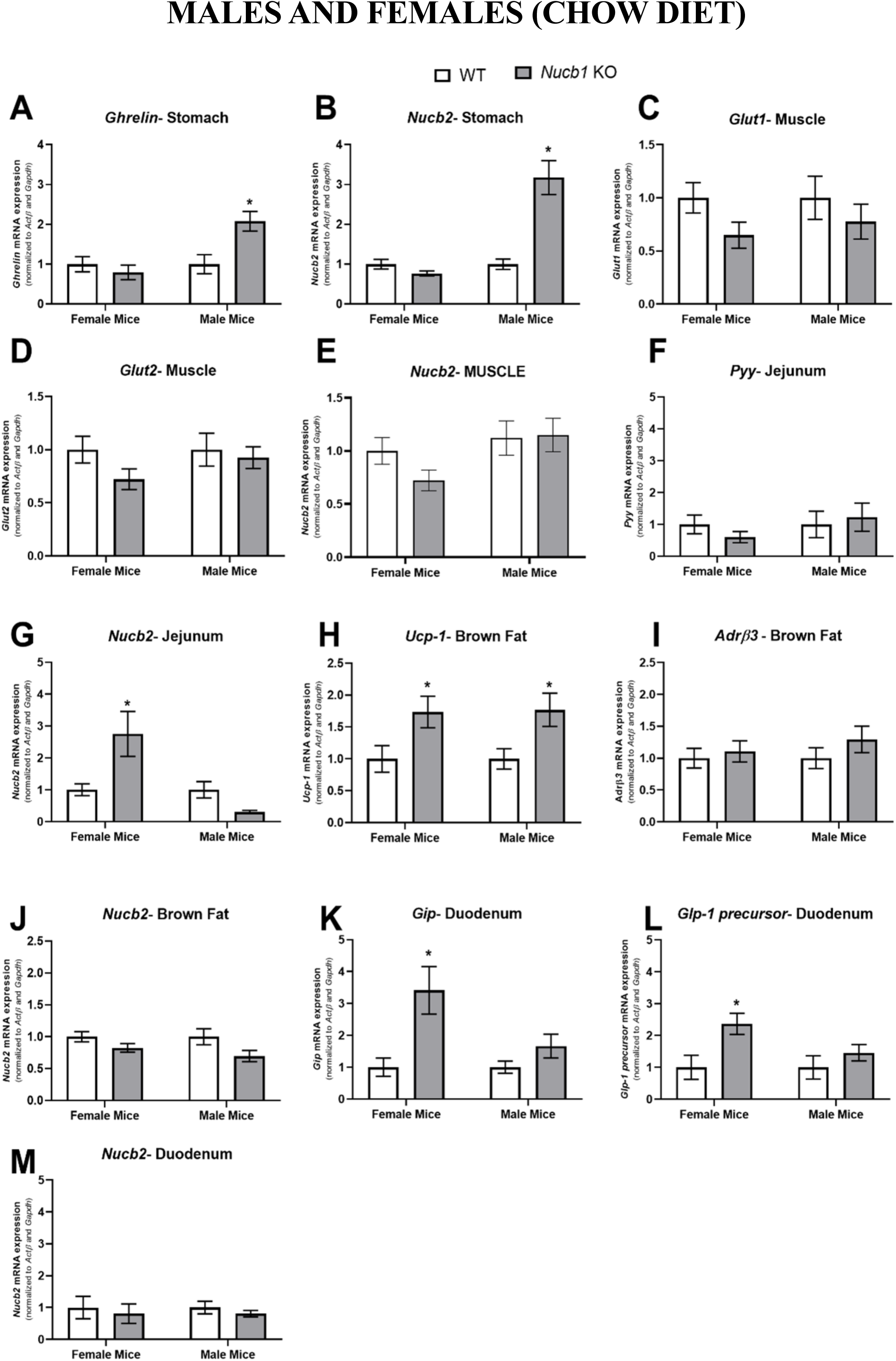
Metabolic gene expression of tissues obtained from *Nucb1* KO mice fed chow diet. The number of samples: >8 tissues/ group/sex. Asterisks show significant differences between WT and KO groups.

## 4. Discussion

Our previous research identified biological roles for NUCB1-derived NLP, including feeding suppression and energy metabolism regulation in male rats [14], insulinotropic effects in MIN6 cells [10], and an anorectic effect in goldfish [11]. NUCB1/NLP co-localizes insulin in the pancreatic islet beta cells of mice [13]. The first goal of our research was to determine if the disruption of *Nucb1* affects whole-body energy homeostasis in mice. The disruption of *Nucb1* caused an increase in food intake in female mice fed a chow diet, but not in male mice, over a 24 h period. Chronic administration of NLP using osmotic minipumps over a 7-day period decreased day 3 and day 4 food intake, with normalization of feeding starting to appear from day 4 dark phase [14]. In the research presented here, while short-term adverse effects were found, the genetic disruption of *Nucb1* did not increase food intake and bodyweight gain in mice over a longer term (weekly monitoring for 5-6 months). This is likely due to adaptive responses to compensate for the absence of NUCB1 and normalize energy homeostasis, via changes including the upregulation of *Nucb2*. NESF-1, processed from NUCB2 that has a high analogy to NUCB1, exhibits an anorectic effect in rodents and modulates central and peripheral orexigenic and anorexigenic peptides [12]. NLP too affected metabolic regulatory hormone mRNA abundance in rats [14]. It is possible that the disruption of *Nucb1* influence the endocrine milieu, and this might indirectly affect energy balance.

In one report [20], *Nucb2* expression was not affected by age and/or diet in 3-month-old versus 6-month-old WT mice fed a high fat. In the same study, the ablation of *Nucb2* did not affect food intake or bodyweight in mice [20]. Although these results agree with our findings on weekly food intake monitoring, their short-term measurement of food intake for 72 h and one-time measurement of bodyweight (10-month-old mice) were different from our approaches, including continuous long-term monitoring. Moreover, several additional criteria, including two-way mixed ANOVA and GEE statistical analysis were employed here, which showed that the genetic disruption of *Nucb1* indeed affected food intake in a time-, diet- and sex-specific manner in high-fat-fed mice. Overall, the results from a very careful characterization with several factors included for consideration, the *Nucb1* disrupted mice exhibits mild perturbations in feeding.

The disruption of *Nucb1* in male and female mice did not alter basal blood glucose levels in weekly monitoring. Although solely based on *in vitro* studies, NLP was reported to stimulate the secretion of insulin in MIN6 cells [10]. Based on this, we expected an elevation in blood glucose due to the loss of positive effects of NLP on insulin synthesis and secretion from islet beta cells. However, our results did not support this expected outcome. The development of compensatory physiological responses, as mentioned earlier, might have contributed to this, as well. Incretin hormones, including GLP-1 and GIP, are secreted by enteroendocrine cells in the small intestine in response to ingested carbohydrates and are then responsible for the stimulation of post-prandial insulin secretion [21]. Consistently, the level of incretin hormones at mRNA levels were upregulated in *Nucb1* disrupted mice. OGTT and IPGTT are reference methods to assess how oral or intraperitoneal delivery of glucose is handled by the organism. OGTT factors in the role of intestinal incretin hormones. Opposite to what we expected, oral glucose handling was better in male *Nucb1* KO mice fed chow diet, while no changes were not observed in female KO mice. Similarly, the disruption of *Nucb1* improved IP glucose handling in male mice fed control and high-fat diet. An increase in incretin hormone transcript abundance (*Gip* and *Glp-1 precursor*) was found in female KO mice fed a chow diet, suggesting a possible role for incretins in the better handling of glucose in *Nucb1* disrupted mice. Another reason for better normalization of glucose following oral or intraperitoneal delivery is an increase in glucose uptake in the target tissues (especially muscle, liver, and adipose tissue) [22] and changes in insulin sensitivity because of genetic disruption of *Nucb1*. Besides, our lab has previously reported that the NESF-1-treated rats had no effects on blood glucose levels, while it increased insulin, and decreased glucagon in rats during an OGTT [23]. A positive effect for NESF-1 on gluconeogenesis in the liver was proposed in that study based on gluconeogenic enzyme mRNA changes, as well as NESF-1 caused an inhibition of glucose uptake in muscle [23]. If NESF-1 and NLP have similar positive effects in increasing glucose in circulation, then the disruption of *Nucb1* would elicit the opposite effects. In fact, our findings suggest that the genetic disruption of *Nucb1* improves insulin sensitivity, and this could be due to a decrease in gluconeogenesis and an increase in glucose uptake. Comprehensive GEE analysis demonstrated a significant general effect of group on blood glucose handling during IPGTT for control and high-fat-fed mice, which supports our findings. Overall, GEE analysis of OGTT and IPGTT results showed that *Nucb1* KO mice fed high-fat diet handled blood glucose better in a sex-, diet- and time point-specific manner. Consistent with our findings, global *Nucb2* deficient male mice fed high-fat diet normalized circulating blood glucose through the increase of hepatic glucose uptake during glucose infusion to maintain euglycemia [20]. Considering the similarity between NESF-1 and NLP, the same effect could occur in *Nucb1* disrupted mice when fed high-fat diet.

To determine whole-body energy homeostasis in *Nucb1* disrupted mice over a 24 h period, CLAMS was used. An increase in food intake during the 24 h period in *Nucb1* disrupted mice was found, but this was observed only in selected cohorts (chow-fed males and females and 10% fat diet-fed male mice). This is a clear indication of how important nutrient composition is in such studies. An increase in food intake of male *Nucb1* KO mice was associated with an increase in preproghrelin (orexigen) mRNA abundance, which might be a contributing factor in the increased food intake of male mice. Our unpublished results show that NESF-1 suppresses preproghrelin mRNA in stomach endocrine cells. It is possible that NLP too regulates different short-term appetite regulatory signals to elicits it effects in maintaining energy balance. This was indeed evident in our sub-cutaneous infusion study in rats, where a number of metabolic regulators mRNAs were found altered []. However, different compensatory physiological responses in the multiple redundant milieu that regulates metabolism might have developed over a longer period due to the genetic disruption of this peptide in mice. This could be a reason why we did not observe major overall changes in the longer-term data in mice of both sexes.

Besides feeding, NUCB1/NLP and NUCB2/NESF-1 are implicated in the control of water intake. Subcutaneous infusion of NLP over 7 days reduced water intake in male rats [14]. Meanwhile, intracerebroventricular (ICV) administration of NESF-1 decreased water intake in rats [24]. This anti-dipsogenic effect was observed under water restriction or after a hypertonic challenge. Interestingly, the anti-dipsogenic effect appeared to be dissociated from the anorexigenic effect of NESF-1, as a higher dose of injected ICV was required for the reduction of water intake. However, the injection of a lower dose of NESF-1 into the subfornical organ, a brain structure that regulates fluid homeostasis, decreased food intake and potently increased water intake in rats [24]. Our finding extends these observations that in the genetic disruption of *Nucb1*, the cumulative water intake was significantly different among female groups at different time points of the dark phase, being higher for KO mice fed chow diet. Since no differences were observed during the light phase, no general group effect was detected (p = 0.2705), but a significant interaction between factors was found (p < 0.0001). The impact on water intake due to *Nucb1* disruption is sexually dimorphic as no significant differences were observed in cumulative water intake of male mice. The diet composition might have affected water intake as no significant changes was detected in WT and Nucb1 KO mice of both sexes fed high fat diet.

The subcutaneous infusion of NLP in male rats over 7 days decreased total physical activity [14]. We observed that the genetic disruption of *Nucb1* induced a significant increase in physical activity in male mice during the dark phase compared to WT littermates fed a chow diet and 10% fat diet. In contrast, physical activity of female mice fed the same diets was opposite to what was found in male mice in the dark phase. This is yet another sexually dimorphic effect for the genetic disruption of *Nucb1*. The increased dark-phase physical activity might be due to the compensatory response of NESF-1 in the genetic disruption of *Nucb1*. In fact, one study reported that central administration of NESF-1 can cause hyperactivity which recovers to the normal level [25]. In agreement with this finding, our lab previously found that long-term treatment of NESF-1 resulted in increased physical activity on the seventh day of continuous infusion in rats [26]. NESF-1 expression was reported to be stimulated by alpha-melanocyte-stimulating hormone, which triggers behaviours related to anxiety and fear, and possibly anxiety-mediated locomotor activity [27]. Considering the observed mild phenotype in *Nucb1* KO mice, possibly due to the compensatory response by NUCB2/NESF-1 and/or other factors, this potential explanation supports our finding. However, this finding seems to be sexually dimorphic for the genetic disruption of *Nucb*.

Energy balance is regulated by many factors that forms the endocrine and non-endocrine milieu [28]. Previous studies reported that NLP and NESF-1 modulate whole-body energy hemostasis in rats [14, 30]. Male *Nucb1* KO mice fed chow diet exhibited higher energy expenditure (heat) than WT littermates in both light and dark phases, but an increase in energy expenditure in female *Nucb1* KO mice applied only in the light phase suggesting energy balance was affected at time-specific manner. This increase was associated with an increase in *Ucp-1* mRNA level in the brown fat of these mice. NLP infusion in rats upregulated *Ucp-1* abundance and increased thermogenesis which might be a reason for the elevation of energy expenditure. However, the increased energy expenditure in *Nucb1* disrupted mice could be at least partly attributed to the compensatory upregulation of NESF-1 that has a stimulatory role on *Ucp-1* expression. The substrate utilization for energy expenditure was also investigated in this research. The infusion of NLP in rats reduced energy derived from fat while carbohydrate catabolism was unaffected for 7 days [14]. Consistent with previous observations, NESF-1-treated rats used fat as a primary source of energy [29]. While these are pharmacological studies based on testing a single dose of the peptide or one mode of administration, direct comparisons of such studies and genetic loss are not always precise. However, these findings provide further evidence to support the fact that both NLP and NESF-1 are fat-influencing peptides, and during their absence, energy derived from fat might alter. The results presented here demonstrate that the catabolism of neither CHO nor fat was altered during the light phase in both sexes fed chow diet, and the use of CHO by KO male and female mice was increased in the dark phase while lipid catabolism was reduced compared to WT mice. The catabolism of CHO decreased in male *Nucb1* KO mice but enhanced in female KO fed high-fat diet mice during both phases. This suggests that the substrate selection for energy production depends on sex and diet ingredients, and the overall compensatory changes due to *Nucb1* disruption affect substrate utilization in these mice. It is also important to note that observed phenotype for *Nucb1* gene disruption may also be dependent on the species studied.

Metabolic tissue weights were also measured in this research. Although HSI was not different between groups (i.e., WT and *Nucb1* KO) in both sexes for all diets, WAT was higher in *Nucb1* KO mice in both sexes fed chow diet and in male KO mice fed high-fat diet. NUCB2/NESF-1 was earlier reported to suppress adipocyte differentiation in the mouse preadipocyte cells (3T3-L1 cells) [30]. NESF-1 has been known as a novel-depot-specific adipokine that has a role in reducing fat mass [31]. Consistently our lab previously reported that the expression of leptin mRNA was reduced in WAT of NLP-treated rats, while the level of adiponectin and resistin mRNA were elevated [14]. These results support our previous findings that NLP is a fat-influencing protein in sex- and diet-specific manner. While some changes in tissue weights were found in selected groups of mice, no major differences in bodyweight were detected in *Nucb1* disrupted mice in comparison to WT litters. Since the pattern of bodyweight gain was different in 60% fat fed male *Nucb1* KO mice versus WT littermates, a significant interaction between male mice and time was observed in bodyweight gain (p <0.0001). Together, we did not find any major changes in bodyweight due to the disruption of *Nucb1*.

## 5. Conclusions

The results of this research provide some supportive evidence for the importance of endogenous *Nucb1*. While it is not critical for growth and reproduction, a mild metabolic phenotype was observed. It is also acknowledged that these effects were found even in the presence of possible compensatory changes in NUCB2/NESF-1, which has structural and functional similarities with *Nucb1*. Another point to note is that the phenotype observed is significant compared to the null phenotype found when more traditional endocrine regulators of metabolism are disrupted or removed [32]. An important finding of our work lies in the interactions observed and highlights the importance of considering the sex, diet and other parameters when interpreting and concluding on research results from KO mice. The reasons for sexual dimorphism found in this research remain unknown, and it clearly is an important topic for follow-up research. The reasons for sexual dimorphism in the mice are also very interesting, and future studies should consider some lines of research in this field.

## 6. Acknowledgments

The authors thank Dr. Sarah Parker, Statistician, WCVM for her kind assistance with our study design and statistical analysis.

## 8. Data availability

The data of this study are available from the corresponding author upon reasonable request.

## 9. Supplementary material

Supplementary information are available as separate document.

## 10. Role of co-authors

AN and E.J.V. planned and performed the experiments, analyzed the data, and prepared the manuscript. J.J.R helped in performing some experiments using high-fat-fed mice. AH established and optimized genotyping technique. SU provided the original ideas, funding for this research, helped design experiments and interpretation of the results, and manuscript review and editing. All authors read and approved the final manuscript.

