## Supplementary material for "Nucleobindin-1 *(Nucb1)* disruption affects feeding, metabolism, and glucose homeostasis in mice in an age-, sex-, diet- and light cycle-dependent manner": Figures

**Table 1**. The ingredients of 10% and 60% fat diets from Research Diets Inc.

**Table 2.** The list of primer forward and reverse sequences and their annealing temperature

**Table 3.** The GEE analysis outcomes of chow-fed mice experiments

**Table 4.** The GEE analysis outcomes for control and high fat fed mice experiments

**Table 1**. The ingredients of 10% and 60% fat diets from Research Diets Inc.

| **Description** | **Ingredients** | **Grams** | **Ingredients** | **Grams** |
|---|---|---|---|---|
| | 60% fat diet | | 10% fat diet | |
| Protein | Casein, Lactic, 30 Mesh | 200.00 g | Casein, Lactic, 30 Mesh | 200.00 g |
| | Cystine, L | 3.00 g | Cystine, L | 3.00 g |
| Carbohydrate | Lodex 10 | 125.00 g | Sucrose, Fine Granulated | 354.00 g |
| | Sucrose, Fine Granulated | 72.80 g | Starch, Corn | 315.00 g |
| | | | Lodex 10 | 35.00 g |
| Fiber | Solka Floc, FCC200 | 50.00 g | Solka Floc, FCC200 | 50.00 g |
| Fat | Lard | 245.00 g | Soybean Oil, USP | 25.00 g |
| | Soybean Oil, USP | 25.00 g | Lard | 20.00 g |
| Mineral | S10026B | 50.00 g | S10026B | 50.00 g |
| Vitamin | Choline Bitartrate | 2.00 g | Choline Bitartrate | 2.00 g |
| | V10001C | 1.00 g | V10001C | 1.00 g |
| Dye | Dye, Blue FD&C #1, Alum. Lake 35-42% | 0.05 g | Yellow FD&C #5, Alum. Lake 35-42% | 0.05 g |
| | Total: | 773.85 g | Total: | 1055.05 g |

**Table 2.** The list of primer forward and reverse sequences and their annealing temperature

| Gene | Accession no. | Primer sequence (5′−3′) | | Temp (°C) |
|---|---|---|---|---|
| | | Forward | Reverse | |
| MS-*Ghrl* | NM_001286404.1 | TCCAAGAAGCCACCAGCTAA | ACAGCTTGATGCCAACATCG | 60 |
| MS-*Gip* | NM_008119.2 | ACAAAGAGGCACAGGAGAGC | AGCCAAGCAAGCTAAGGTCA | 56.2 |
| MS-*Glp-1 precursor* | AF276754.1 | AATCTTGCCACCAGGGACTT | AGTGACTGGCACGAGATGTT | 58 |
| MS-*Pyy* | NM_145435.1 | TTCAGGCCAGAAGGTTTGGA | ACACCGAGATATGAAGTGCCC | 60.6 |
| MS-*Ucp-1* | AH002110.2 | GGCCTCTACGACTCAGTCCA | TAAGCCGGCTGAGATCTTGT | 60 |
| MS-*Adrβ3* | XR_003947220.2 | GCCTTCAACCCGGTCATCTAC | CCTGGGTTCCCGAAGAAGGG | 61 |
| MS-G*lut2* | NM_031197.2 | ATCGCTCCAACCACACTCAG | GCTGAGGCCAGCAATCTGAC | 61 |
| MS-*Glut1* | XM_006502908.2 | CTGCTCATCAACCGCAAC | CTTCTTCTCCCGCATCATCT | 58 |
| MS-*Nucb2* | XR_004934144.1 | AACACGAGCGGAGAGAGTAT | AGGGTCCAATCCATCAGTCT | 60 |
| MS-*Actβ* | BC138614 | CCACTGCCGCATCCTCCTCC | CTCGTTGCCAATAGTGATGAC | 60 |
| MS-*Gapdh* | XM_036165840 | GACATCAAGAAGGTGGTG | ATACCAGGAAATGAGCTTGACAAA | 59 |

**Table 3.** The GEE analysis outcomes of chow-fed mice experiments

| Test | Parameters | Run No. | Model: Significant main effects of factors/predictors on parameters | Model: Significant interactions factors/predictors on parameters |
|---|---|---|---|---|
| Weekly Monitoring | Bodyweight | 1 | Sex, Age | |
| | | 2 | | Sex X Group |
| | | 3 | | Sex X Group X Age |
| | Blood Glucose | 1 | Sex | |
| | | 2 | | Sex X Group, Group X Age |
| | | 3 | | Sex X Group X Age |
| | Food Intake | 1 | Sex, Age | Sex X Group, Group X Age |
| | | 2 | | Sex X Group X Age |
| | | 3 | | |
| OGTT | Blood Glucose | 1 | Sex, Time point | |
| | | 2 | | Sex X Group |
| | | 3 | | Sex X Group X Time point |
| IPGTT | Blood Glucose | 1 | Time point | |
| | | 2 | | Sex X Group |
| | | 3 | | Sex X Group X Time point |
| CLAMS Experiment | Food Intake | 1 | Mice Group, Day phase | |
| | | 2 | | Sex X Group |
| | | 3 | | Sex X Group X Day phase |
| | Water Intake | 1 | Sex, Day phase | |
| | | 2 | | Sex X Group |
| | | 3 | | Sex X Group X Day phase |
| | Heat | 1 | Day phase | |
| | | 2 | | Sex X Group |
| | | 3 | | Sex X Group X Day phase |
| | RER | 1 | Day phase | |
| | | 2 | | Sex X Group, Group X Day phase |
| | | 3 | | Sex X Group X Day phase |
| | Total physical activity | 1 | Sex, Day phase | |
| | | 2 | | Sex X Group |
| | | 3 | | Sex X Group X Day phase |

**Table 4.** The GEE analysis outcomes for control and high fat fed mice experiments

| | Parameters | Run No. | Model: Significant main effects of factors/predictors on parameters | Model: Significant interactions factors/predictors on parameters |
|---|---|---|---|---|
| Weekly Monitoring | Bodyweight | 1 | Sex, Age, Diet | |
| | | 2 | | Sex X Diet, Age X Diet, Sex X Age |
| | | 3 | | Sex X Group X Age, Sex X Group X Diet |
| | | 4 | | Sex X Group X Age X Diet |
| | Blood Glucose | 1 | Group, Sex, Diet | |
| | | 2 | | Sex X Group, Group X Age, Sex X Diet |
| | | 3 | | Sex X Group X Age, Sex X Group X Diet |
| | | 4 | | Sex X Group X Age X Diet |
| | Food Intake | 1 | Sex, Age, Diet | |
| | | 2 | | Sex X Diet, Age X Diet, Sex X Age |
| | | 3 | | Sex X Group X Age, Sex X Group X Diet |
| | | 4 | | Sex X Group X Age X Diet |
| OGTT | Blood Glucose | 1 | Sex, Time point, Diet | |
| | | 2 | | Sex X Group, Group X Diet, Group X Time point |
| | | 3 | | Sex X Group X Time point, Sex X Group X Diet |
| | | 4 | | Sex X Group X Time point X Diet |
| IPGTT | Blood Glucose | 1 | Group, Time point, Sex, Diet | |
| | | 2 | | Sex X Group |
| | | 3 | | Sex X Group X Time point, Sex X Group X Diet |
| | | 4 | | Sex X Group X Time point X Diet |

…. to be continued in the next page …

| | | | Model | |
|---|---|---|---|---|
| | Parameters | Run No. | Significant main effects of factors/predictors on parameters | Significant interactions factors/predictors on parameters |
| CLAMS Experiment | Food Intake | 1 | Day phase, Sex, Diet | |
| | | 2 | | Group X Diet |
| | | 3 | | Group X Day phase X Diet |
| | | 4 | | Sex X Group X Day phase X Diet |
| | Water Intake | 1 | Day phase, Sex, Diet | |
| | | 2 | | Sex X Group, Group X Day phase, Group X Diet |
| | | 3 | | Sex X Group X Day phase, Group X Diet X Day phase |
| | | 4 | | Sex X Group X Day phase X Diet |
| | Heat | 1 | Day phase, Sex, Diet | |
| | | 2 | | Sex X Group, Group X Day phase, Group X Diet |
| | | 3 | | Sex X Group X Day phase, Sex X Group X Diet |
| | | 4 | | Sex X Group X Day phase X Diet |
| | RER | 1 | Day phase, Sex, Diet | |
| | | 2 | | Sex X Group, Group X Diet |
| | | 3 | | Sex X Group X Diet |
| | | 4 | | Sex X Group X Day phase X Diet |
| | Total physical activity | 1 | Sex | |
| | | 2 | | Sex X Group |
| | | 3 | | Sex X Group X Day phase, Sex X Group X Diet |
| | | 4 | | Sex X Group X Day phase X Diet |
